# Spatial expression programs of the intestinal follicle-associated epithelium

**DOI:** 10.1101/2021.04.06.438564

**Authors:** Noam Cohen, Hassan Massalha, Shani Ben-Moshe, Adi Egozi, Milena Rozenberg, Keren Bahar Halpern, Shalev Itzkovitz

## Abstract

The intestine is lined with isolated lymphoid follicles (ILFs) that facilitate sampling of luminal antigens to elicit immune responses. Technical challenges related to the scarcity and small sizes of ILFs and their follicle-associated epithelium (FAE) impeded the characterization of their spatial gene expression programs. Here, we combined RNA sequencing of laser capture microdissected tissues with single molecule transcript imaging to obtain a spatial gene expression map of the ILF and its associated FAE in the mouse small intestine. We identified zonated expression programs in both follicles and FAEs, with a decrease in enterocyte anti-microbial and absorption programs and a partial induction of expression programs normally observed at the villus tip. We further identified Lepr+ sub-epithelial telocytes at the FAE top, which are distinct from villus-tip Lgr5+ telocytes. Our analysis exposes the epithelial and mesenchymal cell states associated with ILFs.

## Introduction

The intestine is lined with mucosa-associated lymphoid follicles that promote homeostatic response against luminal microbiota [1]. These lymphoid follicles include 7-10 large Peyer’s patches (1-2 mm in diameter) [2], as well as smaller solitary intestinal lymphoid tissues (SILT) that are up to 200 µm in diameter. SILTs consist of a spectrum of developed tissues, ranging from cryptopatches (CP) to mature isolated lymphoid follicles (ILF), numbering at around 100-200 along mouse small intestine [3,4]. Peyer’s patches and mature ILFs are covered by follicle-associated epithelium (FAE), forming a dome-like shape [5]. The FAE is mainly composed of enterocytes, with a minority of interspersed Microfold (M) cells. Secretory goblet cells have been shown to be scarce along the Peyer’s patch FAE, resulting in a thinner mucus layer at the luminal surface, and by that facilitating close interactions with microbial antigens [6]. The thin mucous layer enables apical-basolateral transport of antigens from the lumen across the FAE into the lamina propria by M cells, mediating the mucosal immune response [7]. This transport program is different from the normal function of the intestinal epithelium, which consists of selective transport of nutrients with a simultaneous blocking of microbial access to the surface.

Several studies described the transcriptomes of FAE cells and M cells in Peyer’s patches [8–10], however technical limitations related to the smaller sizes of ILFs impeded their similar transcriptomic characterization. Here we applied laser capture microdissection RNA sequencing (LCM RNA-seq [11,12]) and single molecule fluorescence in-situ hybridization (smFISH, [13,14]) to generate a spatial expression map of ILFs and their associated FAE in the mouse small intestine. We identify zonated expression programs in ILFs and FAE, with a decrease in anti-microbial and absorption programs at the FAE top and a compensatory increase in secreted anti-microbial peptides at the FAE boundaries. The FAE top consists of enterocytes that partially induce genes normally observed at the villus tip, and are in contact with Lepr+ sub-epithelial telocyte cells.

## Results

### Spatial mapping of ILF and FAE gene expression

To investigate the spatial heterogeneity of epithelial, immune and stromal gene expression in the ILF and its associated FAE, we used LCM to isolate five segments. These included three epithelial segments consisting of FAE top (FAET), FAE bottom (FAEB), and an adjacent segment at the bottom of a neighboring villus (denoted Villus bottom, VB), and two immune segments consisting of the ILF top (ILFT) and ILF bottom (ILFB) (Fig 1A-C). We used mcSCRBseq [15,16], a sensitive UMI-based protocol to sequence the RNA of the dissected fragments. We identified distinct zonated expression and enriched gene sets in each of the segment (Fig 1D, Fig S1). We used these LCMseq expression programs as a basis for extensive smFISH validations, to establish their statistical power.

**Fig 1.**
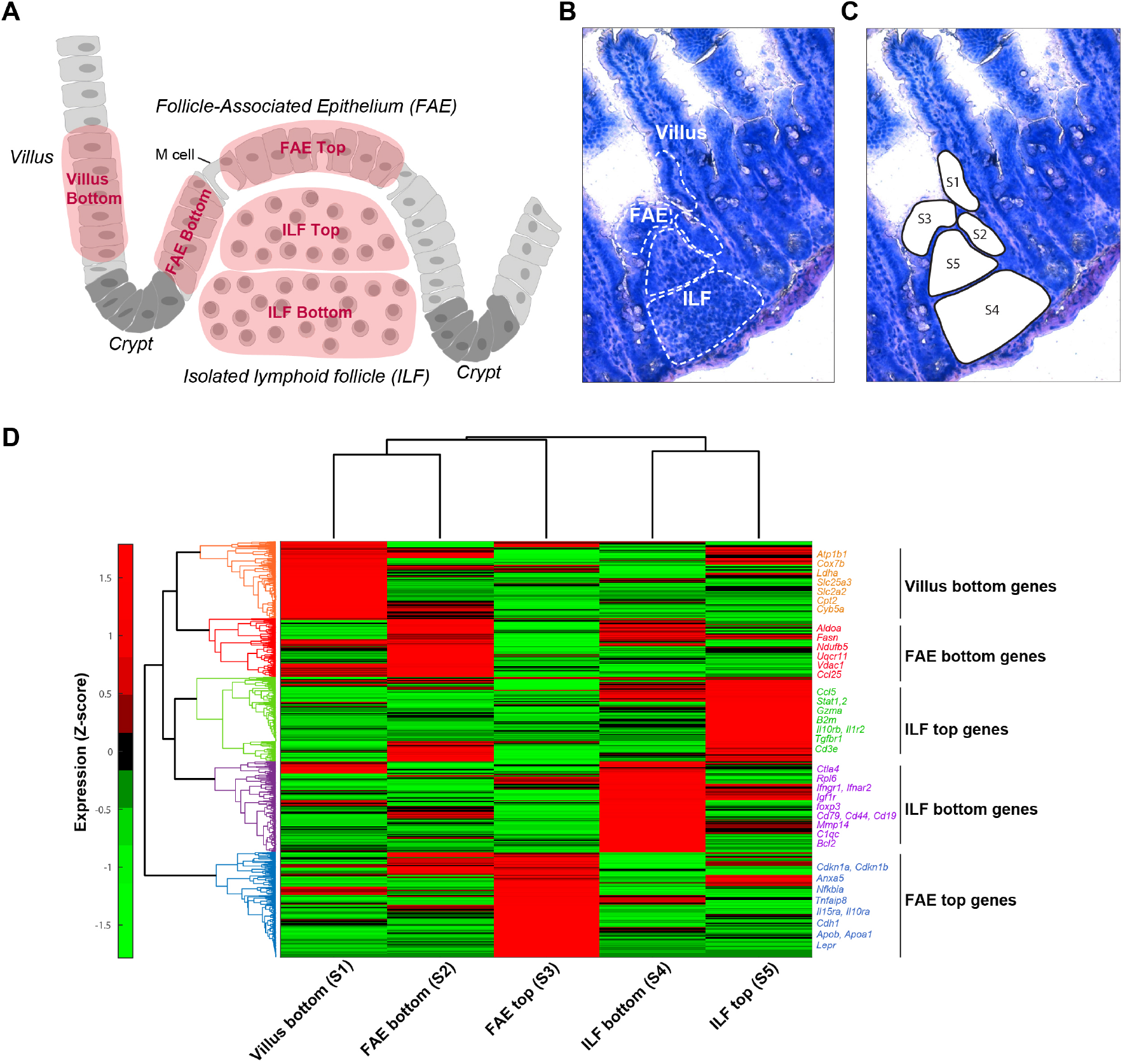
LCM RNA-seq of follicle-associated epithelium (FAE) and isolated lymphoid follicle (ILF). (**A**) An illustration of small intestinal region including FAE, ILF, adjacent villus and crypts, showing the five dissected segments in red: Villus bottom (S1), FAE bottom (S2), FAE top (S3), ILF bottom (S4) and ILF top (S5). (**B-C**) A bright field microscopy image (20x magnification) of FAE, ILF and the adjacent villus before and after laser dissection of the five segments (S1-S5). (**D**) Clustergram of LCM RNA-seq data showing gene mean expression Z-score of the five segments (S1-S5). Selected genes with high expression are shown on the right side of the clustergram for each cluster, colored according to the cluster color.

We first examined the spatial expression programs at the ILF compartment. We identified higher expression of different B-cell markers at the bottom of the ILF such as *Cd19* (Figs S2A,B, S3A,D). In contrast, T cell markers did not show a spatial bias towards the ILF bottom (Fig S2C,D). Rather, using smFISH we found that *Cd3e*, a classic T cell marker, and *Gzma*, a marker of cytotoxic CD8+ T cell subset were highly abundant at the ILF top, infiltrating into the FAE top layer (Fig S3B,C,E,F). This spatial pattern of lymphocytes in the ILF resembles the architecture previously observed in Peyer’s patches, which exhibit a core of B cells and a mantle of T cells [17].

The FAE segments exhibited elevated expression of M cell markers, including *Anxa5* (Fig S3G-I) that were higher in the FAE top compared to FAE bottom and villus bottom. Yet M cells constituted only 8.8% ±1.7% of all FAE cells (mean of five different smFISH images, Fig S3I). We next turned to examine the zonated properties of the remaining >90% epithelial cells in the FAE.

### Anti-microbial and absorption programs are downregulated in FAE top

The mucosal layer that lines the intestinal epithelium is a critical barrier against bacteria. This barrier consists of a mucus layer that contain antimicrobial peptides. Previous in-situ imaging studies on FAE of the large Peyer’s patches demonstrated a lower abundance of the mucus-secreting goblet cells [18,19], as well as a decline in enterocyte anti-microbial genes [6]. Our spatial analysis enabled global characterization of these and other zonated antimicrobial programs with high spatial resolution in FAE associated to the smaller ILFs.

Our LCMseq showed decreased levels of Muc2 in the FAE (Fig 2A). Using smFISH we validated the lower abundance of Muc2+ goblet cells in the FAE with 3.6%±1.0% compared to 9.3%±1.1% in the adjacent villus (mean of five different smFISH images, Fig 2A, B). We also found a decline in the epithelial expression of *Pigr*, encoding the polymeric immunoglobulin receptor that facilitates transcytosis of IgA antibodies from the lamina propria to the lumen [6] (Fig 2A, C, I). *Nlrp6*, encoding a component of the microbial sensing inflammasome [20] also exhibited a steady decline from the villus bottom to the FAE top (Fig 2A, E, I). Unlike *Nlrp6, Muc2* and *Pigr*, which decreased in expression in both the FAE bottom and top compared to the adjacent villus bottom, we found that *Reg3g*, encoding a defensin secreted antimicrobial peptide, decreased at the FAE top, but exhibited an increase in the FAE bottom, compared to the adjacent villus bottom (Fig 2A, D, I). The decline in antimicrobial programs at the FAE top may contribute to the required enrichment for bacteria at this sampling zone, whereas the compensatory higher expression at the FAE boundaries may help to prevent microbial access into the neighboring crypts. Notably, genes encoding ribosomal proteins exhibited similar spatial expression patterns as *Reg3g*, with a decrease in the FAE top and a compensatory increase at the FAE bottom compared to an adjacent villus bottom (Fig S4A). This increased expression of ribosomal proteins at the FAE boundaries could potentially facilitate the elevated translation of the secreted defensin proteins in this zone.

**Fig 2.**
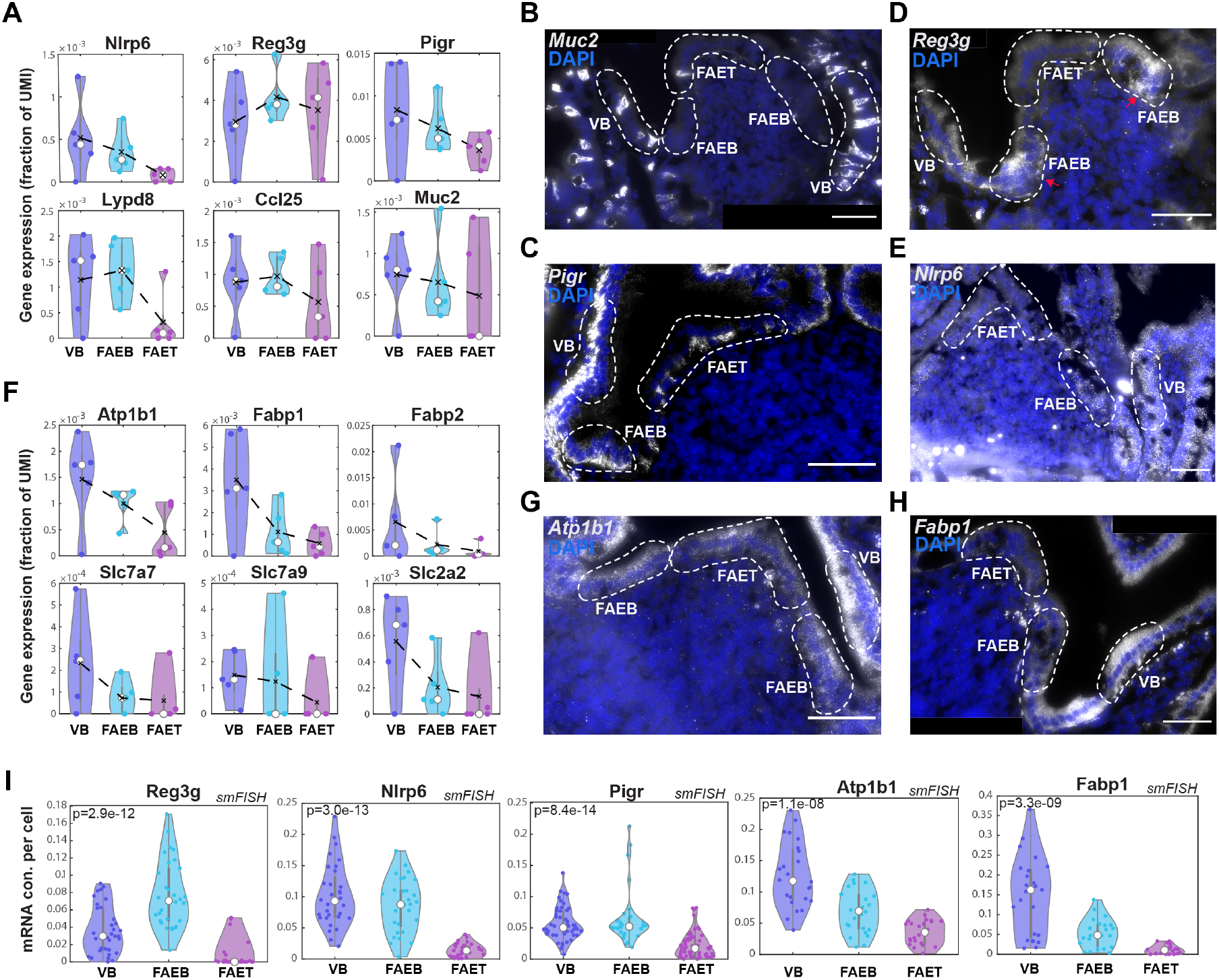
Downregulation in antimicrobial and absorption expression programs. (**A**,**F**) Violin plots of LCMseq gene expression (fraction of UMI) showing downregulated expression of anti-microbial genes (**A)** - *Nlrp6, Reg3g, Pigr, Lypd8, Ccl25* and *Muc2* and nutrient absorption genes (**F)** - *Atp1b1, Fabp1, Fabp2, Slc7a7, Slc7a9* and *Slc2a2* in FAE top (FAET) compared to FAE bottom (FAEB) and villus bottom (VB). Black dashed lines represent mean values, white dots represent median values of five repeats of at least 3 individual FAE from two mice. (**B-E, G-H**) Representative smFISH images showing decreased expression of *Muc2* (**B**), *Pigr* (**C**), *Reg3g* (**D**), *Nlrp6* (**E**), as well as, *Atp1b1* (**G**) and *Fabp1* (**H**) in FAET compared to FAEB and VB. *Reg3g* reduced expression in FAET is compensated by higher expression in FAEB, marked with red arrows. White dashed lines delimit FAET, FAEB and VB areas. DAPI staining for cell nucleus in blue. Scale bar - 50 µm. (**I**) Violin plots of dot quantifications of smFISH of *Reg3g, Nlrp6* and *Pigr*, as well as, *Atp1b1* and *Fabp1* (at least 10 cells were segmented per zone in a pool of 3-5 individual FAE from at least 2 mice).

Along with the reduction of antimicrobial programs, our analysis further revealed a steady decrease in key genes associated with nutrient absorption towards the FAE top. These included the sodium-coupled transporter *Atp1b1*, the fatty acid binding proteins *Fabp1, Fabp*2, as well as different solute carrier genes, such as the glucose transporter *Slc2a2*, and the amino-acid transporters *Slc7a7, Slc7a9* (Fig 2F-I and Fig S4B).

### FAE top enterocytes exhibit a partial villus tip program

Previous reconstruction of spatial gene expression patterns along the intestinal villus axis revealed a substantial up-regulation of dozens of genes in enterocytes at the villus tip [11]. These included cell-adhesion genes such as *Cdh1*, encoding E-cadherin, apolipoproteins, stress-associated transcription factors such as *Jun, Fos* and *Klf4*, as well as the purine-metabolism immune-modulatory genes *Ada, Nt5e* and *Slc28a2* (Fig 3A). We asked whether FAE top enterocytes exhibit similar elevation of these genes. We found that some villus tip enterocyte genes, such as *Cdh1, Cdkn1a* (encoding the protein P21), and the apolipoprotein genes *Apoa1* and *Apoa4* were elevated at the FAE top. In contrast, *Fos, Jun, Klf4* and *Slc28a2* were not elevated at the FAE top (Fig 3B-K). The increase in Apoa proteins, which are essential components of chylomicrons used to transport absorbed lipids to the body, is surprising, given that the absorption machinery was largely downregulated in the FAE top zone (Fig 2F, S4B). This increase could be related to additional anti-inflammatory roles of apolipoproteins [21,22]. Our analysis therefore demonstrates that FAE top enterocytes exhibit a partial induction of the expression program normally observed at the villus tip.

**Fig 3.**
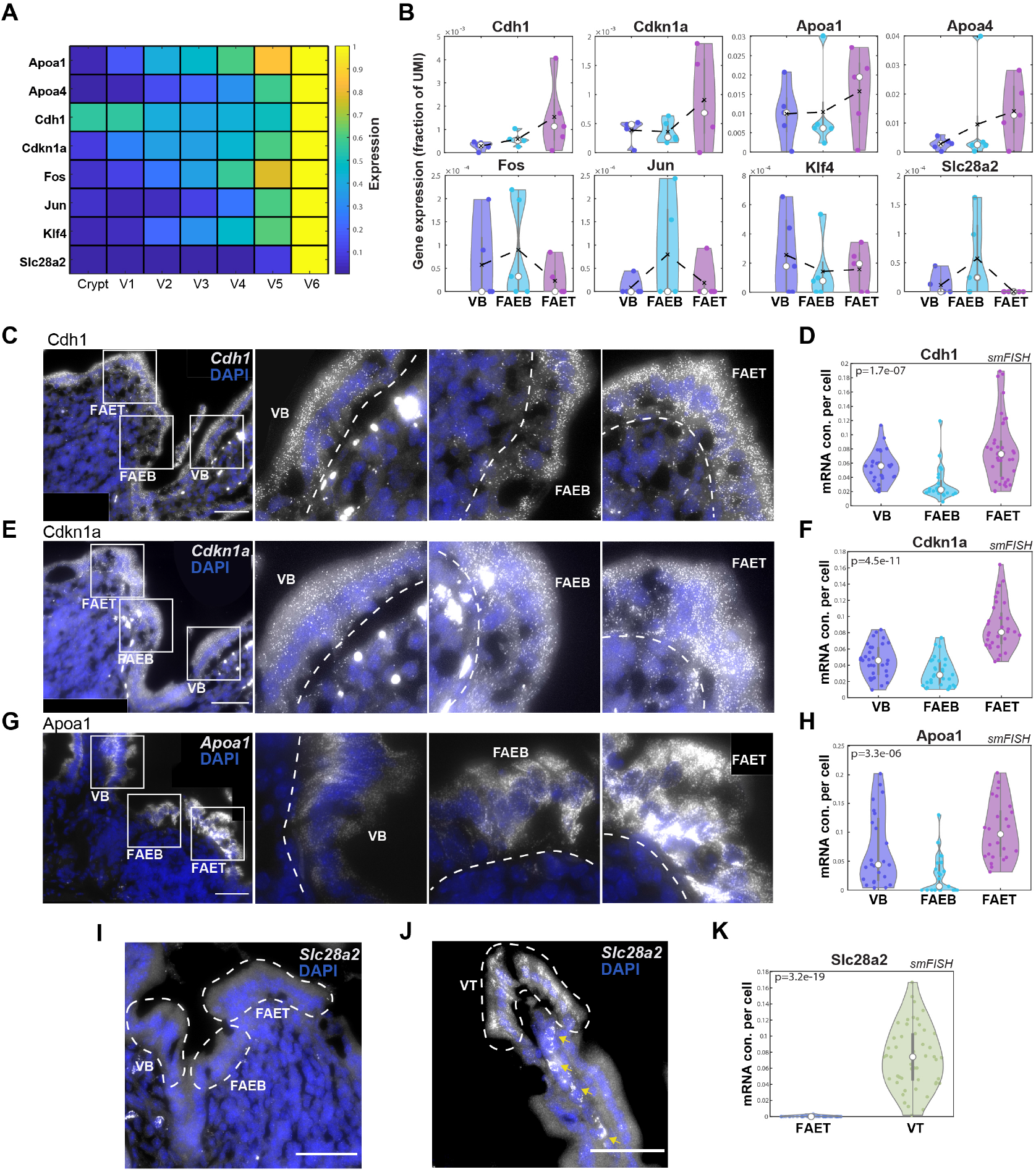
FAE top enterocytes exhibit a partial villus tip program. (**A**) Heatmap of epithelial gene expression along the crypt-villus axis (V1-villus bottom and V6-villus top), showing upregulated expression of tip program genes (data from [11]). (**B**) Violin plots of gene expression (fraction of UMI) showing increased expression of *Cdh1, Cdkn1a, Apoa1* and *Apoa4*, but not *Fos, Jun, Klf4* and *Slc28a2* at the FAE top. Black dashed lines represent mean values and white dots represent median values of five repeats of at least three individual FAE from two mice. (**C, E, G**) Representative smFISH images showing upregulated expression of *Cdh1* (**C**), *Cdkn1a* (**E**) and *Apoa1* (**G**) in the FAET compared to FAEB and VB. In each panel, left image shows the ILF/FAE, followed by magnifications of the labelled white-boxed regions. Scale bar is 50 µm. DAPI staining of cell nucleus in blue. White dashed lines delimit FAET, FAEB and VB areas. (**D, F, H**) Violin plots of dot quantifications of smFISH experiments. P values of Kruskal Wallis tests are presented. (**I-J**) Representative smFISH images showing no expression of *Slc28a2* in FAET, FAEB and VB, and high expression at the villus tip (VT). Yellow arrows point at autofluorescent blobs. (**K**) Violin plot of smFISH dot quantification of *Slc28a2* expression in FAET compared to VT. The smFISH validations (**D, F, H, K**) showing the concentration (con.) of dots (mRNA molecules) per cell area (at least 10 cells were segmented per zone in a pool of 3-5 individual ILF/FAEs from at least two mice).

### FAE top telocytes are marked by Lepr and are distinct from villus tip telocytes

PDGFRα+ sub-epithelial telocytes are slender elongated cells that line the intestinal epithelium and have recently been shown to be important niche cells that shape zonated epithelial expression programs [23,24]. Villus tip telocytes, marked by the gene *Lgr5*, are regulators of the villus tip enterocyte program [25]. We argued that our LCM RNA-seq of the epithelial segments may also include sub-epithelial telocytes, in addition to epithelial cells, due to their adjacency to the epithelial layer. To identify potential telocyte gene expression, we extracted 1,765 genes that were previously shown to be substantially more highly expressed in telocytes compared to enterocytes (Methods). Among these potential telocyte marker genes, we found elevated expression at the FAE top of *Lepr*, encoding the leptin receptor (Fig 4A-B). *Lepr* was previously shown to be elevated in crypt mesenchymal cells in the colon [26]. Using smFISH combined with immunostaining for PDGFRα, we validated that FAE top telocytes express significantly higher levels of *Lepr* compared to the FAE bottom and adjacent villus bottom (Fig 4C, F). In contrast, *Lgr5* was highly expressed in villus tip telocytes (as well as in the crypt epithelial stem cells), but was undetectable in the sub-epithelial layers of the FAE (Fig 4C-E). Our analysis therefore identifies *Lepr+* sub-epithelial telocytes distinct from the *Lgr5+* telocytes at the villus tip.

**Fig 4.**
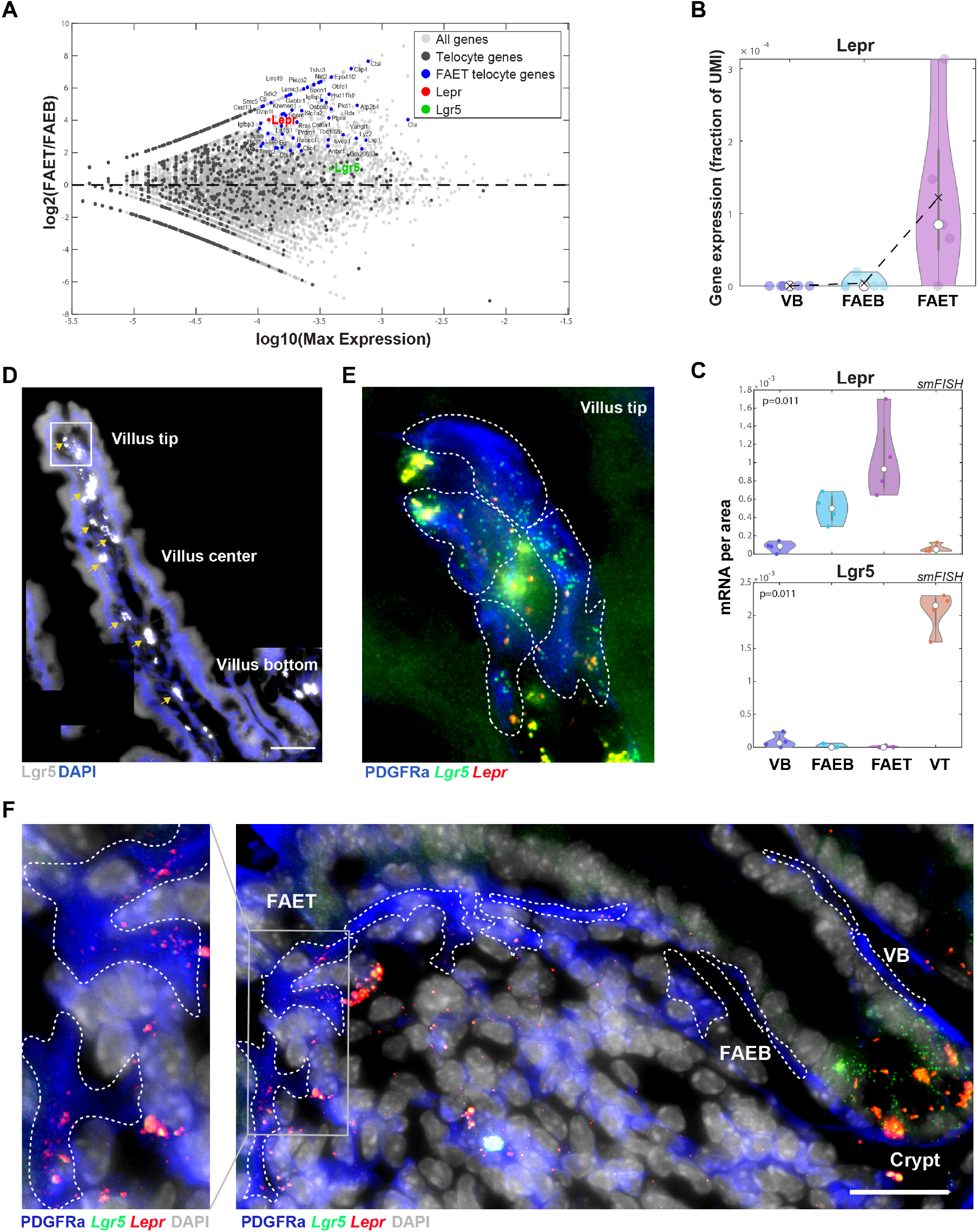
Lepr+ telocytes are zonated to FAE top. (**A**) MA plot showing log2 of the expression ratios between FAET and FAEB vs. log10 of the max-expression. Gray dots – all genes, dark gray dots - telocyte genes, blue dots - highly expressed telocyte genes upregulated in FAET. *Lepr* gene marked in red and *Lgr5* gene in green (expression curated from previous scRNAseq papers, see Methods). (**B**) Violin plot showing high expression of *Lepr* (as a fraction of UMI) in FAET. Black dashed lines represent mean values and white dots represent median values of five repeats of at least three individual FAE from two mice. (**C)** Violin plots of quantification of smFISH experiments for *Lepr* and *Lgr5* showing mRNA concentrations (con.) per zone in smFISH images (four repeats of different zones). P values of Kruskal Wallis tests are presented. (**D-F)** smFISH validations of *Lgr5* and *Lepr* expression in villus tip and FAET. (**D**) smFISH image showing *Lgr5* expression in gray of a full length villus (tip, center and bottom). Yellow arrows point at autofluorescent blobs. Scale bar-50 µm. (**E**) Blow up of villus tip showing *Lgr5* expression (green) but not *Lepr* expression (red) in PDGFRα telocytes. **(F**) smFISH image showing *Lepr* expression in gray in FAE. Scale bar-25 µm. Blow up showing *Lepr* expression (red) in PDGFRα telocytes (blue) in FAET but not in FAEB and VB. *Lgr5* expression is absent from FAET telocytes and observed in the crypt stem cells. White dashed lines delimit PDGFRα telocytes.

## Discussion

ILFs, along with Peyer’s patches, are critical sites of intestinal immune surveillance, yet their scarcity and small sizes render them hard to isolate. Our analysis provides a comprehensive spatial atlas of gene expression of ILFs and their associated epithelium. We found that FAE enterocytes exhibit a gene expression signature that is distinct from both enterocytes at the villus bottom and at the villus tip. FAE top enterocytes express lower levels of antimicrobial genes, as well as lower levels of the nutrient absorption genes. These features seem to be tuned towards maintaining a microenvironment that is optimal for efficient sampling of bacterial antigens by M cells and immune cells, rather than nutrient absorption.

What are the factors that could elicit the distinct gene expression programs at the FAE top? FAE enterocytes are localized at around the same physical distance from the crypt as villus bottom enterocytes. Moreover, previous work has shown that FAE eneterocytes are continuously migrating and shedding from the FAE tip, similarly to villus enterocytes [27]. Yet, the FAE operates under a unique microenvironment compared to the villus bottom. Luminal bacterial concentrations at the FAE tip should be higher due to the reduced mucous layer and reduced secretion of anti-microbial peptides, and therefore more similar to the luminal microenvironment at the villus tip [11]. We found that, unlike villus-tip enterocytes, FAE top enterocytes are in contact with *Lepr*+ telocytes, which could provide different niche signals than their villus tip *Lgr5*+ telocytes counterparts. Indeed, the expression of the purine-metabolism immune-modulatory genes *Ada, Nt5e* and *Slc28a2* at the villus tip seems to be controlled by *Lgr5*+ telocytes [25], potentially explaining their reduced expression at the FAE top. Our study forms the basis for the future exploration of the regulatory molecules that shape FAE zonation. It will be interesting to expand our study to cryptopatches and colonic ILFs, as well as to the characterization of ILFs in perturbed states such as germ-free mice and in models of inflammatory diseases.

## Methods

### Animal experiments

All animal studies were approved by the Institutional Animal Care and Use Committee (IACUC) of the Weizmann Institute of Science. C57bl6 male mice at the age of 12-15 weeks (Harlan laboratories) were kept in SPF conditions and fed with regular chow *ad libitum*. LCM experiments were conducted on the Jejunum segments extracted from two WT mice. Tissues were transected on top of Wattman paper soaked in cold PBS and then were immediately embedded without fixation in OCT (Scigen, 4586) on dry ice. Frozen blocks were transferred to -80°C until use. For smFISH experiments, jejunum tissues were harvested from 4 WT mice and fixed in 4% formaldehyde (J.T. Baker, JT2106) for 3 h, incubated overnight with 30% sucrose in 4% formaldehyde, then embedded in OCT in the form of swiss-rolls and were kept at -80°C until use.

### Laser capture microdissection (LCM)

LCM experiments were performed as was previously described [11] with some modifications. Briefly, ten serial sections of 10 µm thickness were attached on polyethylene-naphthalate membrane-coated 518 glass slides (Zeiss, 415190-9081-000), air-dried for 20 s at room temperature, washed in 70% ethanol for 25 s, incubated in water for 25 s (Sigma-Aldrich, W4502), stained with HistoGene Staining Solution for 20 s (ThermoFisher Scientific, KIT0401) and washed again in water for 25 s. Stained sections were dehydrated with subsequent 25 s incubations in 70%, 95%, and 100% EtOH and air-dried for 60 s before microdissection. The experiments were performed using UV laser-based unit (PALM-Microbeam, Zeiss) and a bright field imaging microscope (Observer.Z1, Zeiss). PALM x40 lenses were used to catapult and collect ILF and FAE tissue segments into 0.2 ml adhesive cap tubes (Zeiss, 415190-9191-000). ILFs were identified histologically as about 200 µm width aggregates of immune and stromal cells in association to a dome-like epithelial layer. Our data included 5 repeats (rep1-5) of 5 ILFS/FAE segments, defined as: ILF bottom, ILF top, FAE bottom, FAE top and Villus bottom (S1-S5) (see Fig 1A-C). Each repeat included at least three independent ILF/FAE, with a total area of 50,000 µm^2^ per segment. Rep1-Rep2 were obtained from Mouse #1 and Rep3-Rep5 from Mouse #2. Pooled sections were resuspend in 7 µl lysis buffer: RLT lysis buffer (Qiagene, 1015762) supplemented with 0.04M DTT, immediately transferred to dry ice and stored at −80 °C until the preparation of RNA libraries.

### LCM RNA-seq

LCM samples were washed from lysis buffer by mixing with an equal portion of SPRI bead solution (Beckman Coulter, A63881), incubated 5 min R/T and washed twice with 100% EtOH under magnetic field. RNA captured on bead was elucidated using 10 µl of RT reaction mixture (Thermo Fisher) (1× Maxima H Buffer, 1 mM dNTPs, 2 µM TSO* E5V6NEXT, 7.5% PEG8000, 20U Maxima H enzyme, 1 µl barcoded RT primer) and was taken to RT reaction (42°C, 90 min and inactivation in 80°C, 10 min). The subsequent steps were applied as previously reported in the of cDNA library preparation protocol for mcSCRB-seq [15] with the following modifications. The cDNA was amplified with 15-18 cycles, depending on the cDNA concentration indicated by qRT– PCR quality control. Then, 0.6 ng of the amplified cDNA was converted into the sequencing library with the Nextera XT DNA Library Preparation Kit (Illumina, FC-131-1024), according to the protocol supplied. Quality control of the resulting libraries was performed with a High Sensitivity DNA ScreenTape Analysis system (Agilent Technologies, 5067-5584). Library concentration was determined using the NEBNext Library Quant Kit (Illumina, E7630). Libraries were diluted to a final concentration of 2.8 pM in HT1 buffer (supplemented with the kit) and loaded on 75-cycle high-output flow cells (Illumina, FC-404-2005) and sequenced on a NextSeq 550 (Illumina). Raw and processed sequencing data are available in the GenBank GEO database NCBI (GSE168483).

### Bioinformatic analysis

Illumina output sequencing raw files were converted to FASTQ files using bcl2fastq package. To obtain the UMI counts, FASTQ reads were aligned to the mouse reference genome (GRCm38.84) using zUMI package [28] and STAR with the following parameters: RD1 16 bp, RD2 66 bp with a barcode (i7) length of 8 bp. For UMI table see Table S1. Subsequent analysis was done with Matlab R2018b. UMI table was filtered to retain only protein coding genes, based on GRcm38.84 ensembl acquired via BioMart using biomaRt R package version 2.44.1 (Table S1). Mitochondrial genes were further removed and the remaining UMI counts for each sample were normalized to the sum of UMI counts of all genes that individually took up less than 5% of the UMI count sum in any of the samples. Mean expression was calculated over all samples from the same zone (Table S1). Gene expression was calculated as fraction of UMI counts over the sum of UMIs (Table S1) and was presented in violin plots. Clustering was performed using the Clustergram function in Matlab with correlation distance over all genes with maximal mean expression above 10^−4^. Gene set enrichment analysis was done using MSigDB_Hallmark_2020 database of Enrichr [29,30]. The list of genes in each set is specified in Table S2. For extracting telocyte genes, we used previous datasets of cell-type specific gene expression in the mouse small intestine and compared the mean expression over villus tip and crypt telocytes [25] to the mean expression over zonated crypt-villus populations [11], goblet cells and enteroendocrine cells [8]. Telocyte genes were defined as those with mean expression above 10^−5^ in telocytes and 4-fold higher expression in telocytes compared to epithelial cells. M cell marker genes (Fig S3G) were obtained from previous analyzed data [8]. To identify T cell and B cell markers we analyzed a previous scRNAseq dataset [31]. We calculated the maximal mean expression among cells annotated as CD4 or CD8 and compared it to the mean expression of B cells. We considered genes with expression above 5*10^−4^ in the corresponding cell type and extracted the 20 genes with the highest expression ratio between the two cell types. Kruskal Wallis tests were used to assess statistical significance as p value<0.05.

### Single molecule FISH (smFISH)

The smFISH experiments were conducted as was previously described [13] with some modifications. Briefly, 8µm thick sections of fixed jejunum were sectioned and placed onto poly L-lysine (Sigma-Aldrich, P8920) coated coverslips, then fixed again in 4% FA in PBS for 15 min R/T followed by 70% Ethanol dehydration for 2h in 4°C. Tissues were treated for 10 min R/T with proteinase K (10 µg/ml Ambion AM2546) and washed twice with 2× SSC (Ambion AM9765), then incubated in wash buffer (20% Formamide Ambion AM9342, 2× SSC) for 10 min R/T. Next, the tissues were mounted with the hybridization buffer (10% Dextran sulfate Sigma D8906, 20% Formamide, 1 mg/ml E.coli tRNA, Sigma R1753, 2× SSC, 0.02% BSA, Ambion AM2616, 2 mM Vanadyl-ribonucleoside complex, NEB S1402S) mixed with 1:3000 dilution of specific probe libraries and were transferred to an overnight incubation at 30°C. Probe libraries were designed using the Stellaris FISH Probe Designer Software (Biosearch Technologies, Inc., Petaluma, CA). For probe library list see Table S3. After overnight incubation the tissues were washed with wash buffer supplemented with 50 ng/ml DAPI (Sigma-Aldrich, D9542) for 30 min at 30 °C and washed with GLOX buffer (0.4% Glucose, 1% Tris, 10% SSC). For the detection of telocytes, goat anti PDGFRα primary antibody (AF1062 R&D systems) (8 µg/µl) was added to the smFISH hybridization buffer and Alexa fluor 488 conjugated donkey anti goat as secondary antibody (Jackson laboratories, 705-545-147, 1:400) in GLOX buffer for 20 minutes after staining with DAPI.

Imaging was performed on Nikon eclipse Ti2 inverted fluorescence microscopes equipped with 100x oil-immersion objectives and a Photometrics Prime 95B 25MM EMCCD camera. All images were taken using ×100 magnifications, therefore several fields of view were stitched together to cover the whole ILF/FAE area using Fiji software. Quantification of smFISH was done using ImageM [14]. Three to five imaged tissues from at least two mice were segmented and fluorescent dots were counted and divided by the segmented cell volume to obtain the mRNA concentration per cell. At least 10 cells per region were segmented from 5 different images. Segmentation of ILF top and bottom was done using a border line in the middle of the ILF that was determined based on DAPI staining of cell nucleus and blinded to the smFISH channel. Specifically, segmentation and dot counting of telocytes was done using Fiji based on the immune-fluorescent staining of the surface expressed PDGFRα. Region of interest (ROI) was manually segmented and dots were counted per area of four different images from two mice. Kruskal Wallis tests were used to assess statistical significance as p value<0.05.

---

**Fig S1.**
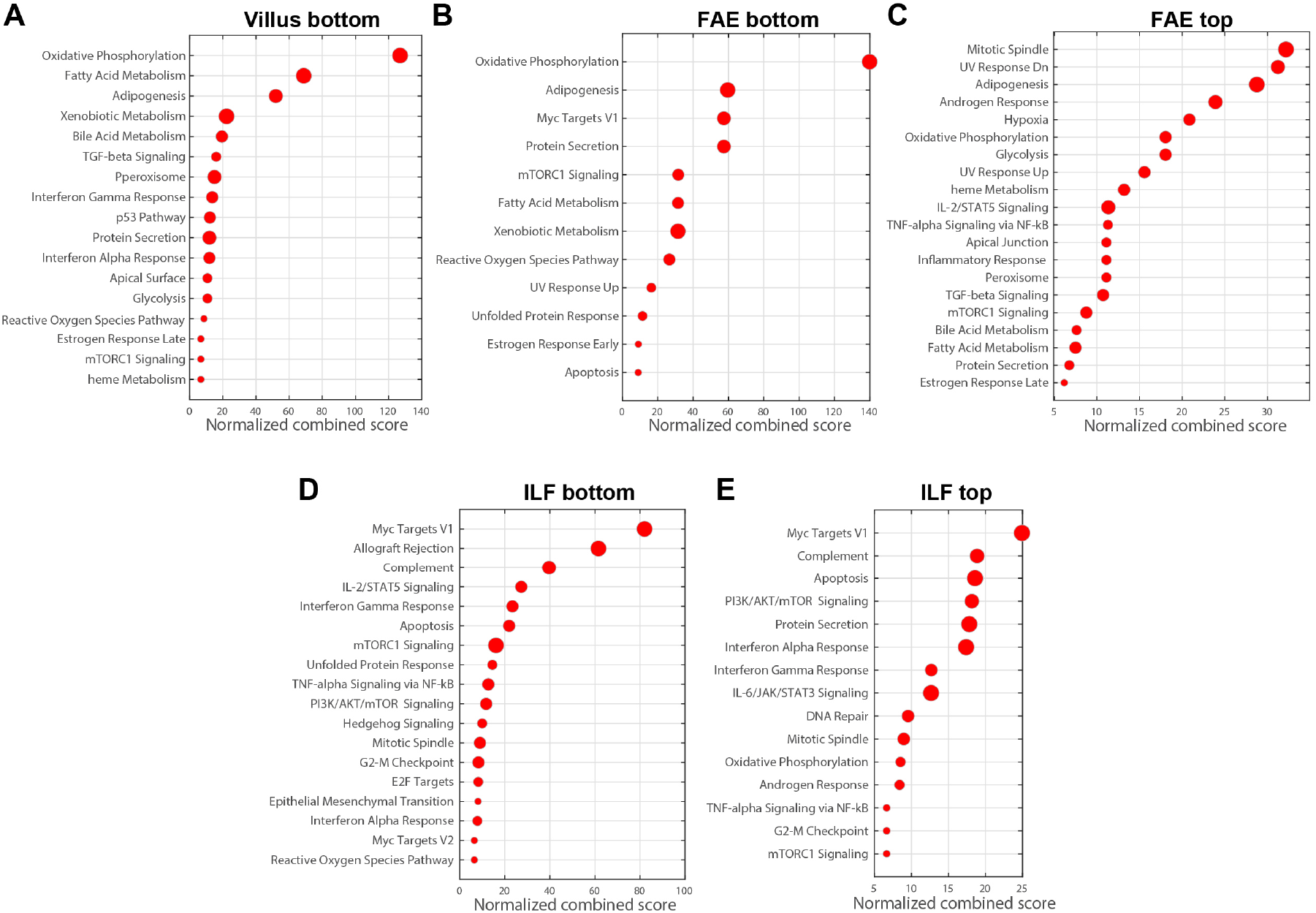
Gene set enrichment. (**A-E**) Dot plots generated by Enrichr (see methods) showing normalized combined score (with q value less than 0.1) of upregulated gene sets for Villus bottom, FAE bottom, FAE top, ILF bottom and ILF top. The size of each red dot is in accordance with the number of genes in each set (for list of genes for each set see Table S2).

**Fig S2.**
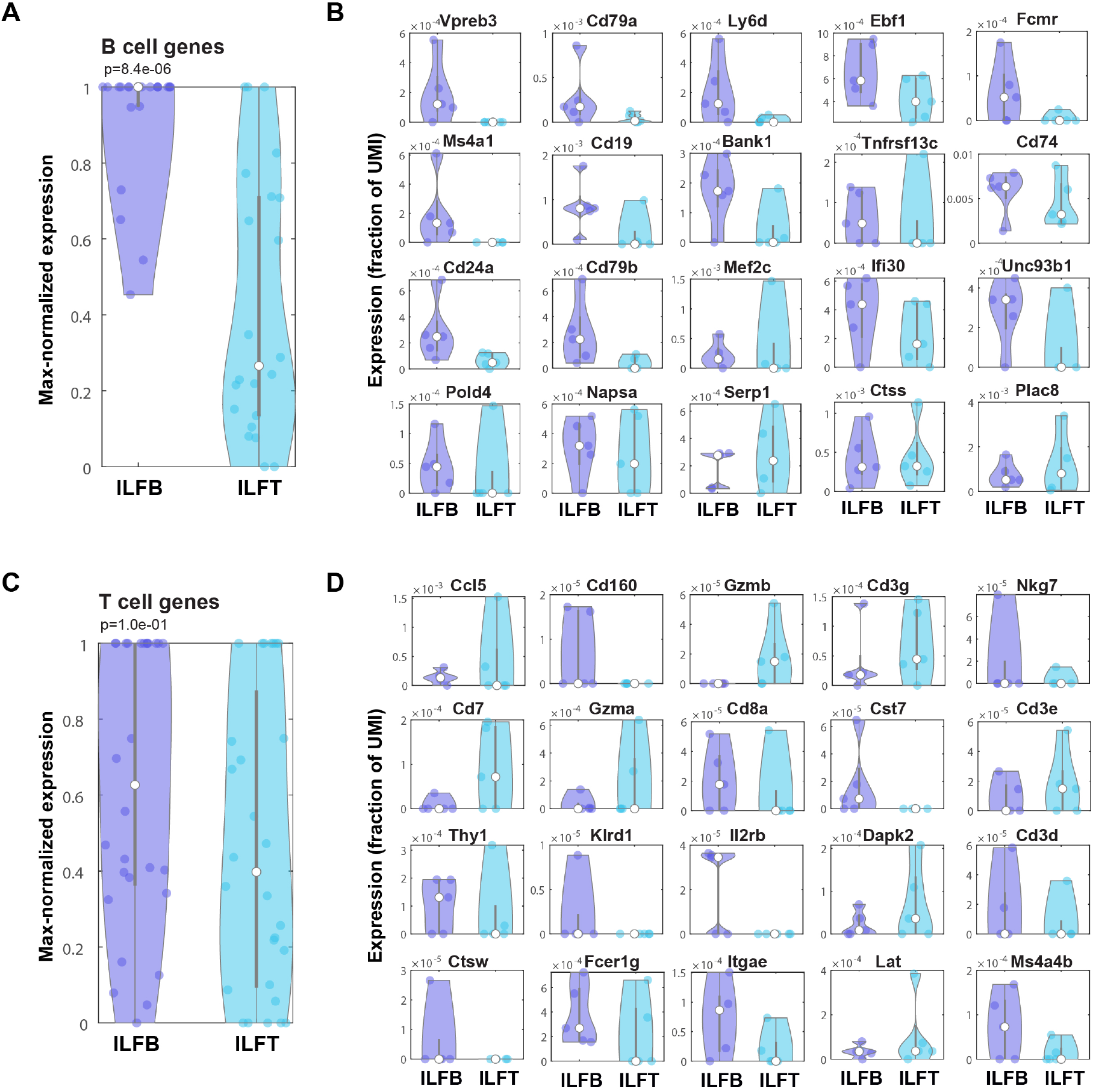
B cells but not T cells are zonated towards the ILF bottom. (**A**,**C**) Violin plots showing max-normalization of B and T cell signature gene expression (methods). B cell markers are significantly zonated to the ILFB, whereas T cell markers exhibited no significant bias to the ILFB. P values of Kruskal Wallis tests are presented. (**B**,**D**) Violin plots showing 20 genes with the highest expression ratio (fraction of UMI) between the B cells (**B)** and T cells (**D)** (Methods). White dots represent median values.

**Fig S3.**
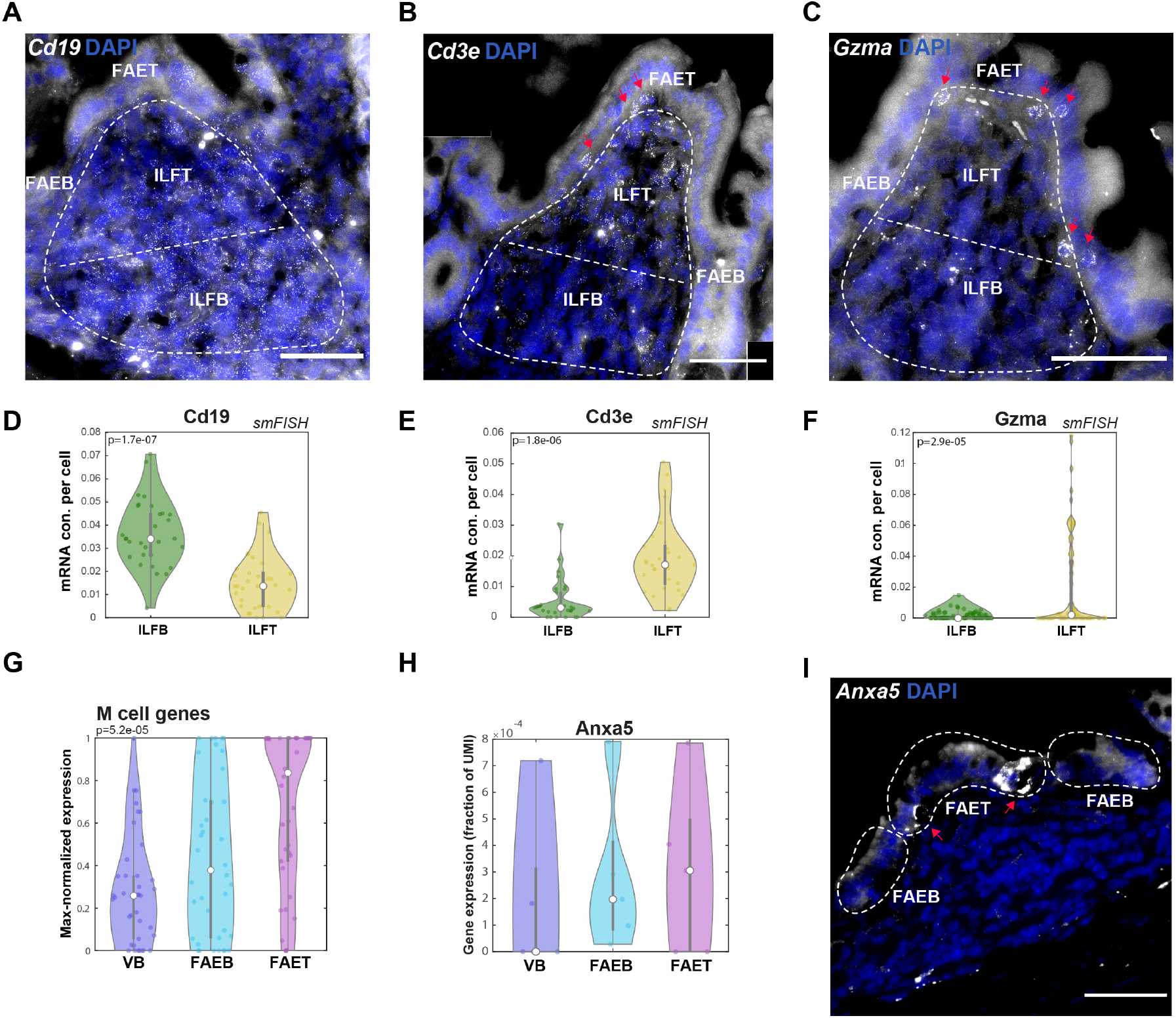
Zonation of immune cell subsets in ILF and M cells in FAE. (**A**,**B**,**C**) smFISH validations showing increased expression of *Cd19* (**A**) *Cd3e* (**B**) and *Gzma* (**C**) at ILF bottom. White dashed lines delimit segment ILF areas and a border line in the middle between ILFT and ILFB. Red arrows mark *Cd3e* and *Gzma* expressing cells penetrating to the FAET layer. DAPI staining for cell nucleus in blue. Scale bar-50 µm. (**D**,**E**,**F**) Violin plots of dot quantifications of smFISH signals of *Cd19, Cd3e* and *Gzma*, showing the concentration (con.) of dots (mRNA molecules) per cell area (at least 10 cells were segmented per zone in a pool of 3-5 individual ILF/FAEs from at least two mice). (**G**) Violin plots for the max-normalized expression of M cell signature genes (methods) showing an upregulation expression of M cell genes in FAET compared to FAEB and VB. (**H**) A violin plot showing upregulation of *Anxa5* expression in FAET compared to FAEB and VB. (**I**) smFISH validation showing *Anxa5* expression in the FAET. White dashed lines delimit FAET and FAEB areas and red arrows mark cells with higher expression levels of *Anxa5*. Scale bar-50 µm. P values of Kruskal Wallis tests are presented.

**Fig S4.**
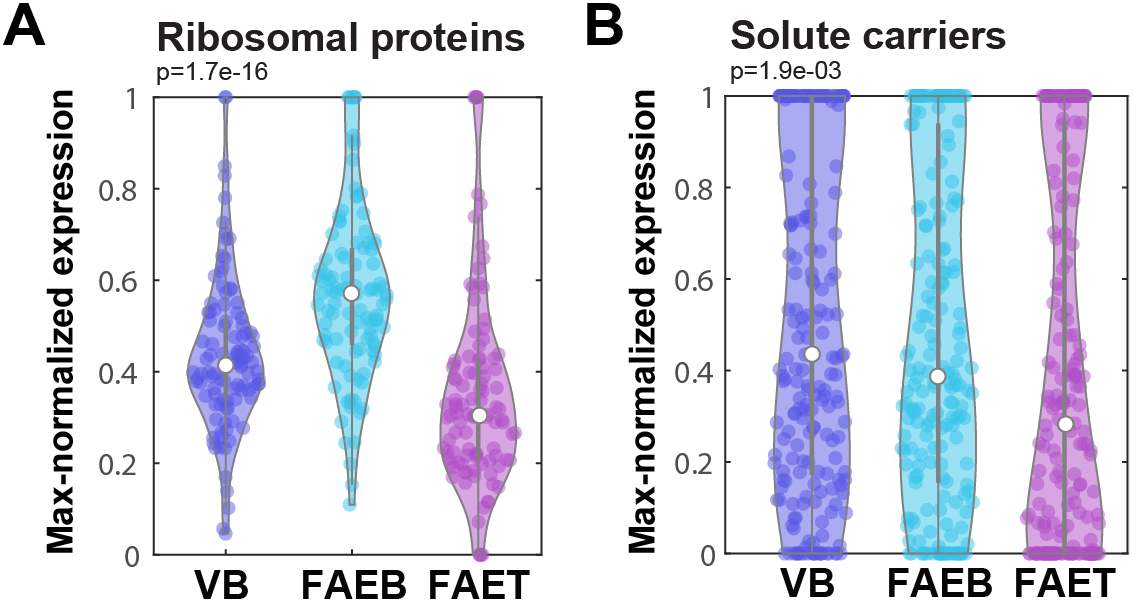
Downregulated expression of ribosomal proteins and solute carriers in FAE top. (**A**,**B**) Violin plots showing downregulated max-normalized expression of ribosomal proteins (**A**) and solute carriers (**B**) in FAET compared to FAEB and VB. P values of Kruskal Wallis tests are presented. White dots represent median values.

**Table S1 - UMI table, Filtered UMI table and mean expression table**. (**Tab1**) UMI table of 24090 genes expressed in S1 (VB), S2 (FAEB), S3 (FAET), S4 (ILFB) and S5 (ILFT) of five experimental repeats obtained from two mice (Mouse#1 and #2) (**Tab2**) UMI table of only protein coding genes (16312 genes). (**Tab3**) Mean expression and standard error of the five experimental repeats.

**Table S2 - Output of enriched set of genes from Enrichr analysis**. Five tabs for each segment: S1 (VB), S2 (FAEB), S3 (FAET), S4 (ILFB) and S5 (ILFT) showing an output of Enrichr analysis (Methods) for upregulated gene sets (Terms), including p values, adjusted p values, odds ratio and combined score. The genes in each set are specified in the last column.

**Table S3 - Probe list**. A list of all probe libraries used for smFISH experiments in this study.

## Acknowledgements

S.I. is supported by the Wolfson Family Charitable Trust, the Edmond de Rothschild Foundations, the Fannie Sherr Fund, the Dr. Beth Rom-Rymer Stem Cell Research Fund, the Minerva Stiftung grant, the Israel Science Foundation grant No. 1486/16, the Broad Institute□Israel Science Foundation grant No. 2615/18, the European Research Council (ERC) under the European Union’s Horizon 2020 research and innovation program grant No. 768956, the Chan Zuckerberg Initiative grant No. CZF2019□002434, the Bert L. and N. Kuggie Vallee Foundation and the Howard Hughes Medical Institute (HHMI) international research scholar award.

## References

1. Nagatake T, Kunisawa J, Kiyono H. Lymphoid Tissues Associated with Gastrointestinal (GI) Mucosa. In: Natsugoe S, editor. Lymph Node Metastasis in Gastrointestinal Cancer. Singapore: Springer; 2019. pp. 111–126. doi:10.1007/978-981-10-4699-5_5

2. Brayden DJ, Jepson MA, Baird AW. Keynote review: Intestinal Peyer’s patch M cells and oral vaccine targeting. Drug Discov Today. 2005;10: 1145–1157. doi:10.1016/S1359-6446(05)03536-1

3. Buettner M, Lochner M. Development and Function of Secondary and Tertiary Lymphoid Organs in the Small Intestine and the Colon. Front Immunol. 2016;7: 342. doi:10.3389/fimmu.2016.00342

4. Hamada H, Hiroi T, Nishiyama Y, Takahashi H, Masunaga Y, Hachimura S, et al. Identification of multiple isolated lymphoid follicles on the antimesenteric wall of the mouse small intestine. J Immunol Baltim Md 1950. 2002;168: 57–64. doi:10.4049/jimmunol.168.1.57

5. Kanaya T, Williams IR, Ohno H. Intestinal M cells: Tireless samplers of enteric microbiota. Traffic Cph Den. 2020;21: 34–44. doi:10.1111/tra.12707

6. Nakamura Y, Kimura S, Hase K. M cell-dependent antigen uptake on follicle-associated epithelium for mucosal immune surveillance. Inflamm Regen. 2018;38: 15. doi:10.1186/s41232-018-0072-y

7. Mabbott NA, Donaldson DS, Ohno H, Williams IR, Mahajan A. Microfold (M) cells: important immunosurveillance posts in the intestinal epithelium. Mucosal Immunol. 2013;6: 666–677. doi:10.1038/mi.2013.30

8. Haber AL, Biton M, Rogel N, Herbst RH, Shekhar K, Smillie C, et al. A single-cell survey of the small intestinal epithelium. Nature. 2017;551: 333–339. doi:10.1038/nature24489

9. Hase K, Ohshima S, Kawano K, Hashimoto N, Matsumoto K, Saito H, et al. Distinct gene expression profiles characterize cellular phenotypes of follicle-associated epithelium and M cells. DNA Res Int J Rapid Publ Rep Genes Genomes. 2005;12: 127–137. doi:10.1093/dnares/12.2.127

10. Terahara K, Yoshida M, Igarashi O, Nochi T, Pontes GS, Hase K, et al. Comprehensive Gene Expression Profiling of Peyer’s Patch M Cells, Villous M-Like Cells, and Intestinal Epithelial Cells. J Immunol. 2008;180: 7840–7846. doi:10.4049/jimmunol.180.12.7840

11. Moor AE, Harnik Y, Ben-Moshe S, Massasa EE, Rozenberg M, Eilam R, et al. Spatial Reconstruction of Single Enterocytes Uncovers Broad Zonation along the Intestinal Villus Axis. Cell. 2018;175: 1156-1167.e15. doi:10.1016/j.cell.2018.08.063

12. Moor AE, Golan M, Massasa EE, Lemze D, Weizman T, Shenhav R, et al. Global mRNA polarization regulates translation efficiency in the intestinal epithelium. Science. 2017;357: 1299–1303. doi:10.1126/science.aan2399

13. Itzkovitz S, Lyubimova A, Blat IC, Maynard M, van Es J, Lees J, et al. Single-molecule transcript counting of stem-cell markers in the mouse intestine. Nat Cell Biol. 2011;14: 106–114. doi:10.1038/ncb2384

14. Lyubimova A, Itzkovitz S, Junker JP, Fan ZP, Wu X, van Oudenaarden A. Single-molecule mRNA detection and counting in mammalian tissue. Nat Protoc. 2013;8: 1743–1758. doi:10.1038/nprot.2013.109

15. Bagnoli JW, Ziegenhain C, Janjic A, Wange LE, Vieth B, Parekh S, et al. Sensitive and powerful single-cell RNA sequencing using mcSCRB-seq. Nat Commun. 2018;9: 2937. doi:10.1038/s41467-018-05347-6

16. Massalha H, Bahar Halpern K, Abu-Gazala S, Jana T, Massasa EE, Moor AE, et al. A single cell atlas of the human liver tumor microenvironment. Mol Syst Biol. 2020;16: e9682. doi:10.15252/msb.20209682

17. Lycke NY, Bemark M. The regulation of gut mucosal IgA B-cell responses: recent developments. Mucosal Immunol. 2017;10: 1361–1374. doi:10.1038/mi.2017.62

18. Ermund A, Gustafsson JK, Hansson GC, Keita ÅV. Mucus Properties and Goblet Cell Quantification in Mouse, Rat and Human Ileal Peyer’s Patches. PLOS ONE. 2013;8: e83688. doi:10.1371/journal.pone.0083688

19. Ohno H. Intestinal M cells. J Biochem (Tokyo). 2016;159: 151–160. doi:10.1093/jb/mvv121

20. Ghimire L, Paudel S, Jin L, Jeyaseelan S. The NLRP6 inflammasome in health and disease. Mucosal Immunol. 2020;13: 388–398. doi:10.1038/s41385-020-0256-z

21. Georgila K, Vyrla D, Drakos E. Apolipoprotein A-I (ApoA-I), Immunity, Inflammation and Cancer. Cancers. 2019;11. doi:10.3390/cancers11081097

22. Yui Y, Aoyama T, Morishita H, Takahashi M, Takatsu Y, Kawai C. Serum prostacyclin stabilizing factor is identical to apolipoprotein A-I (Apo A-I). A novel function of Apo A-I. J Clin Invest. 1988;82: 803–807. doi:10.1172/JCI113682

23. Aoki R, Shoshkes-Carmel M, Gao N, Shin S, May CL, Golson ML, et al. Foxl1-Expressing Mesenchymal Cells Constitute the Intestinal Stem Cell Niche. Cell Mol Gastroenterol Hepatol. 2016;2: 175–188. doi:10.1016/j.jcmgh.2015.12.004

24. Shoshkes-Carmel M, Wang YJ, Wangensteen KJ, Tóth B, Kondo A, Massasa EE, et al. Subepithelial telocytes are an important source of Wnts that supports intestinal crypts. Nature. 2018;557: 242–246. doi:10.1038/s41586-018-0084-4

25. Bahar Halpern K, Massalha H, Zwick RK, Moor AE, Castillo-Azofeifa D, Rozenberg M, et al. Lgr5+ telocytes are a signaling source at the intestinal villus tip. Nat Commun. 2020;11: 1936. doi:10.1038/s41467-020-15714-x

26. Matsumura S, Kurashima Y, Murasaki S, Morimoto M, Arai F, Saito Y, et al. Stratified layer analysis reveals intrinsic leptin stimulates cryptal mesenchymal cells for controlling mucosal inflammation. Sci Rep. 2020;10: 18351. doi:10.1038/s41598-020-75186-3

27. Sierro F, Pringault E, Assman PS, Kraehenbuhl JP, Debard N. Transient expression of M-cell phenotype by enterocyte-like cells of the follicle-associated epithelium of mouse Peyer’s patches. Gastroenterology. 2000;119: 734–743. doi:10.1053/gast.2000.16481

28. Parekh S, Ziegenhain C, Vieth B, Enard W, Hellmann I. zUMIs -A fast and flexible pipeline to process RNA sequencing data with UMIs. GigaScience. 2018;7. doi:10.1093/gigascience/giy059

29. Chen EY, Tan CM, Kou Y, Duan Q, Wang Z, Meirelles GV, et al. Enrichr: interactive and collaborative HTML5 gene list enrichment analysis tool. BMC Bioinformatics. 2013;14: 128. doi:10.1186/1471-2105-14-128

30. Kuleshov MV, Jones MR, Rouillard AD, Fernandez NF, Duan Q, Wang Z, et al. Enrichr: a comprehensive gene set enrichment analysis web server 2016 update. Nucleic Acids Res. 2016;44: W90–97. doi:10.1093/nar/gkw377

31. Biton M, Haber AL, Rogel N, Burgin G, Beyaz S, Schnell A, et al. T Helper Cell Cytokines Modulate Intestinal Stem Cell Renewal and Differentiation. Cell. 2018;175: 1307-1320.e22. doi:10.1016/j.cell.2018.10.008

